# Convergence of case-specific epigenetic alterations identify a confluence of genetic vulnerabilities tied to opioid dependence

**DOI:** 10.1101/2021.06.15.447736

**Authors:** Olivia Corradin, Richard Sallari, An T. Hoang, Bibi S Kassim, Gabriella Ben Hutta, Lizette Cuoto, Bryan C. Quach, Katreya Lovrenert, Cameron Hays, Berkley E. Gryder, Marina Iskhakova, Hannah Cates, Yanwei Song, Cynthia F. Bartels, Dana B. Hancock, Deborah C. Mash, Eric O. Johnson, Schahram Akbarian, Peter C. Scacheri

## Abstract

Opioid dependence is a highly heterogeneous disease driven by a variety of genetic and environmental risk factors which have yet to be fully elucidated. We interrogated the effects of opioid dependence on the brain using ChIP-seq to quantify patterns of H3K27 acetylation in dorsolateral prefrontal cortical neurons isolated from 51 opioid-overdose cases and 51 accidental death controls. Among opioid cases, we observed global hypoacetylation and identified 388 putative enhancers consistently depleted for H3K27ac. Machine learning on H3K27ac patterns predicts case-control status with high accuracy. We focus on case-specific regulatory alterations, revealing 81,399 hypoacetylation events, uncovering vast inter-patient heterogeneity. We developed a strategy to decode this heterogeneity based on convergence analysis, which leveraged promoter-capture Hi-C to identify five genes over-burdened by alterations in their regulatory network or “plexus”: *ASTN2, KCNMA1, DUSP4, GABBR2, ENOX1*. These convergent loci are enriched for opioid use disorder risk genes and heritability for generalized anxiety, number of sexual partners, and years of education. Overall, our multi-pronged approach uncovers neurobiological aspects of opioid dependence and captures genetic and environmental factors perpetuating the opioid epidemic.

## Introduction

Opioid dependence and the rise of opioid overdose deaths have taken a devastating emotional and economic toll on society. This public health crisis has only been exacerbated during the COVID19 pandemic with reports of a 16 to 25% increase in opioid-related overdose deaths in the US (Center for Disease Control, 2020). Uncovering the neurobiological basis of opioid use disorder (OUD) is urgently needed to drive prevention and treatment interventions. However, current GWAS, with over 10,000 cases, have only replicably identified the *OPRM1* gene (Zhou et al., 2020a) (Hancock et al., 2018), making it an especially challenging phenotype to study. Conservative estimates suggest that hundreds of thousands of samples will be needed to uncover a majority of risk loci (Sullivan et al., 2018). Animal models have produced valuable insights into the biology of disease, but the extent to which these will translate to the complexity of human behavioral traits remains unclear. In lieu of larger GWAS sample sizes that are not currently available from human studies, we present a methodological innovation to shortcut discovery of genes involved in the neurobiology of opioid dependence. We further identify novel genes and pathways that point us towards biology relevant for understanding opioid dependence.

Here, we reason that a successful approach would encompass three key features: (1) the accurate measurement of epigenomic activity and connectivity to genes, (2) the importance of making these measurements in the appropriate cellular context and (3) the need to harness heterogeneous, case-specific alterations. We pioneer such an analysis in OUD through the analysis of histone acetylation landscapes in opioid overdose cases vs. accidental sudden death (physiologically normal) controls. We focus on neurons from dorsolateral prefrontal cortex (DLPFC) as a cell type broadly involved in the neural circuitry of opioid and other drug addictions, as well as, the frequently co-occurring disorders of anxiety and impulsivity. Moreover, animal and human studies have drawn a firm link between long-term vulnerability to addiction and cellular and molecular adaptations in the brain’s reward-processing circuitry (Koob and Volkow, 2010), including changes in gene expression programs in the prefrontal cortex and other target areas of the mesocorticolimbic dopamine system (Browne et al., 2020). We therefore reasoned that mapping the epigenome of DLPFC in opioid overdose death cases vs. controls would be the ideal context to discover regulatory loci important in mediating addictive behaviors.

Tissue-specific regulatory elements consistently show a high correlation with GWAS catalog risk loci in phenotypically relevant tissue types (Maurano et al., 2012) (Trynka et al., 2013) (Finucane et al., 2015). We focus our epigenomic assays on H3K27ac, the canonical mark of active regulatory elements. We interrogate neuronal H3K27ac epigenomes of DLPFC from opioid overdose cases compared with controls. Moreover, by analyzing H3K27ac epigenomes in the context of promoter-bound chromosomal conformations, using a promoter-capture Hi-C reference set from the DLPFC (Jung et al., 2019), we are able to connect individual H3K27ac peaks to promoter targets with high confidence.

Previous H3K27ac epigenomic profiling of colon cancer, metastatic osteosarcoma and numerous other types of cancer (Akhtar-Zaidi et al., 2012) (Cohen et al., 2017) (Morrow et al., 2018) has revealed stark tumor-specific alterations in enhancer activity. We termed these VELs (Variable or Variant Enhancer Loci). Unlike these observations from rapidly dividing cells undergoing a constant evolution from normal to malignant, here we study post-mitotic neurons in the human brain, arguably one of the most stable tissue types in the human body. Thus, we do not anticipate dramatic changes in enhancer activities in this setting. This requires a rethinking of our theoretical framework; instead of single clear changes with potent effects on gene expression, we transition to dispersed, subtle effects that cumulatively tweak gene expression. We hypothesize that direct ascertainment of subtle yet converging heterogeneous variants present in the brain epigenome of fatal opioid overdose cases will illuminate the biological and genetic underpinnings of the traits involved in opioid dependence.

## RESULTS

### Identification of homogeneous H3K27ac hypoacetylation events

We generated H3K27ac ChIP-seq data from neuronal nuclei sorted from postmortem DLPFC dissected from 60 confirmed opioid overdose death cases and 58 age-matched, sudden accidental death controls. A total of 51 cases and 51 controls passed stringent quality control (**Methods, Table S1**) and were selected for further analysis (**Figure 1A, S1**). These samples were sequenced to a mean read depth per sample of 112,215,946 (s.d. = 33,175,526), with no difference in read depth between case and control groups (Welsh unpaired two-sample t-test, two-sided P=0.605, n=102 biologically independent samples, read count mean difference ± SEM: 3418882 ± 6593642; 95% confidence interval (CI): -9684412-16522176; t=0.5185, df=88.09) (**Figure S1**). To our knowledge, this represents one of the most extensive analyses of H3K27ac in human neuronal DLPFC performed to date, and the only one focused on opioid dependence. Across the cohort, we detected 655,505 H3K27ac peaks. Linear regression was used to identify differentially acetylated loci in cases relative to controls, controlling for covariates including age, postmortem interval, ancestry and sex. We observed modest but widespread hypoacetylation of H3K27 in cases versus controls, with 388 peaks showing statistically significant differences (P_Bonf._ < 1×10^−7^) (**Figure 1B,C, Table S2**). An exemplar locus linked to target gene DUAL-SPECIFICITY PHOSPHATASE 4 (*DUSP4*) in DLPFC promoter capture Hi-C data, involved in the addiction-relevant GPCR and MAPK signaling pathways (Blackwood et al., 2021; Qiao et al., 2021), is shown in **Figure 1D**. Strikingly, no significant hyperacetylated peaks in cases were found. Of the 388 significant hypoacetylated peaks, 95.9% were located distal (> 2 kb) to transcription start sites and are putative active regulatory elements. Motifs corresponding to AP1 transcription factors, including FOS, ATF3, and JUNB (P<1×10^−15^) were highly enriched at these regulatory elements (**Figure 1E**). We identified putative gene targets of the differentially acetylated loci using available promoter-capture Hi-C data from DLPFC (Jung et al., 2019).

**Figure 1.**
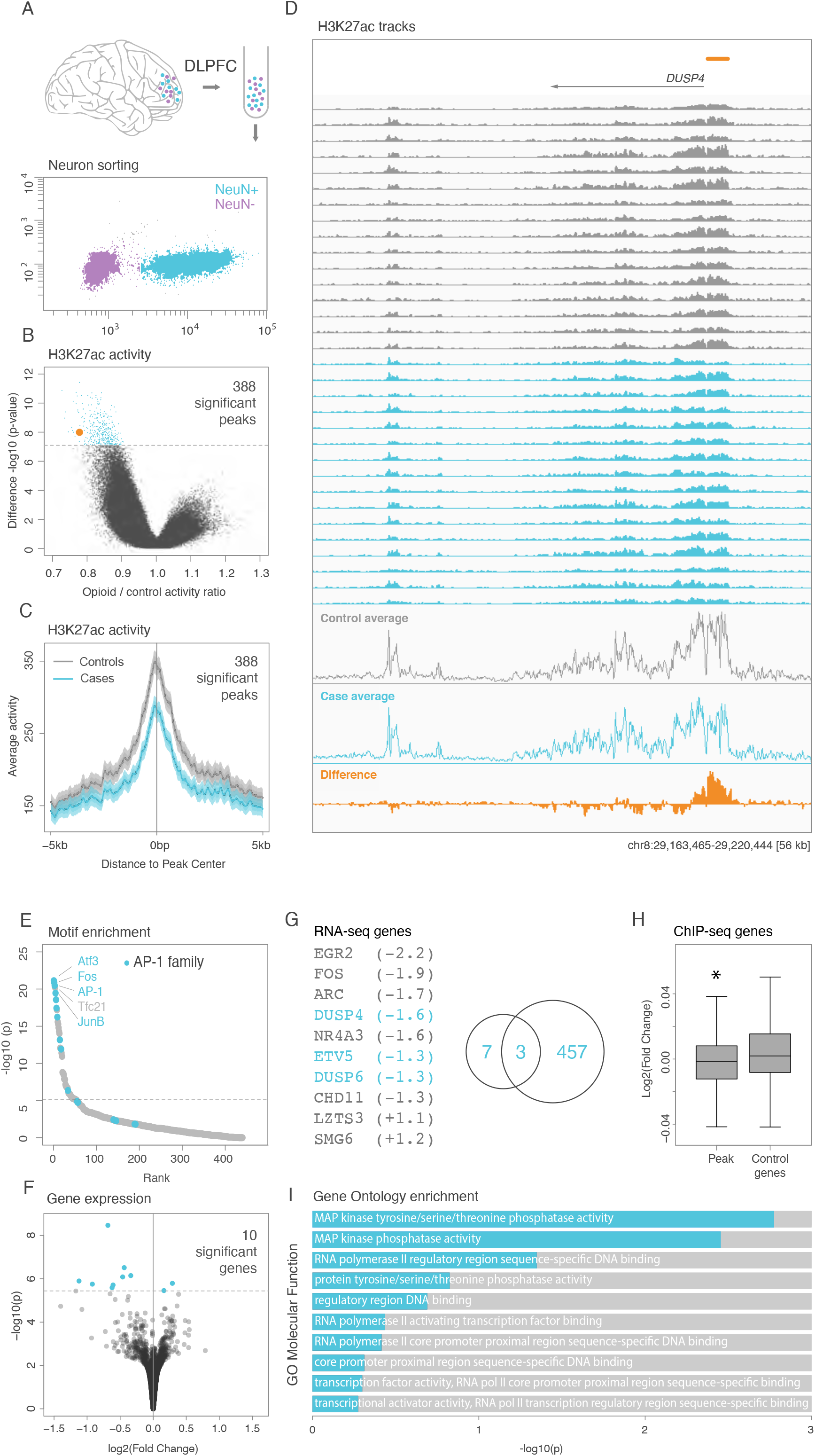
Identification of homogeneous H3K27ac hypoacetylation events. **(A)** Schematic overview of experimental strategy. NeuN^+^ nuclei were identified using FANS from post mortem dorsolateral prefrontal cortex tissues, followed by H3K27ac ChIP-seq. **(B)** Volcano plot of linear regression analysis of ChIP-seq samples. 388 differentially acetylated peaks identified at genome-wide significance are shown in blue. Example locus DUSP4 highlighted in orange corresponding orange bar in D. **(C)** Aggregate H3K27ac ChIP-seq signal across controls (grey) and opioid cases (blue) for 388 differentially acetylated regions. **(D)** Browser image of H3K27ac ChIP-seq results at example locus *DUSP4*. Orange bar highlights differentially acetylated peak. **(E)** Motifs enriched in differentially acetylated peaks ranked by enrichment p-value. **(F)** Volcano plot of differentially expressed genes identified through RNA-seq of bulk tissue samples (n=24 cases and n=27 controls). Bonferroni multi-test correction threshold is shown in dotted line. **(G)** Comparison of differential peaks and genes. (left) Differentially expressed genes with log_2_ fold change (cases/controls) shown in parentheses. Genes in blue overlap gene targets of linear regression peaks. (right) Venn diagram showing 10 differentially expressed genes (left circle) compared to 460 genes linked to linear regression peaks (right circle). **(H)** Tukey boxplot of log_2_ fold-change (cases/controls) in gene expression of gene targets associated with linear regression peaks versus randomly selected genes not associated with a linear regression peak. Welch’s two sample t-test p=0.00067. Outliers not shown. **(I)** Gene ontology enrichment of differentially expressed genes.

We performed RNA-seq analysis of a subset of the cohort (24 cases, 27 controls, bulk DLPFC, not neuronal nuclei sorted) (**Figure 1F)**. Ten genes were identified as differentially expressed, 8 downregulated, 2 upregulated in cases compared to controls (**Figure 1G**, P_Bonf._ < 2.4×10^−6^). The AP1 transcription factor *FOS* was among the significantly reduced genes. Genes assigned by promoter capture Hi-C to the 388 hypoacetylated regulatory elements generally show a modest but statistically significant decrease in gene expression levels in cases compared to controls (P < 0.05) (**Figure 1H**). Overlap of the significant epigenome and transcriptome results highlights 3 genes: *DUSP4, DUSP6, ETV5* (**Figure 1G)**. *DUSP4* and *DUSP6* regulate MAPK-ERK signaling. Consistent with this known biological function, the MAP kinase pathway was the most enriched GO molecular function term associated with the set of differentially expressed genes in opioid cases versus controls (P_adj_< 0.01) (**Figure 1I**) (Chen et al., 2013; Kuleshov et al., 2016). Both the input transcription factors to the 388 loci (FOS AP1 factors) and output MAPK genes are consistent with known addiction pathways (Al-Hasani and Bruchas, 2011) (Blackwood et al., 2021) (Browne et al., 2020) (Qiao et al., 2021) (Brynildsen et al., 2020).

### Seven loci can accurately predict case/control status

The 388 loci identified through linear regression represent only a small proportion of the hundreds of thousands of ChIP-seq peaks evaluated. We therefore applied machine learning (ML) as an agnostic strategy to re-evaluate the entirety of the data. We used Gradient Boosting classification algorithm with ChIP-seq peaks and covariates as potential features to distinguish case/control status (**Methods**) (Chen and Guestrin, 2016). Remarkably, seven ChIP-seq peaks can differentiate opioid cases from controls with high accuracy (5-fold cross-validated mean AUC = 0.972, s.d.=0.037) (**Figure 2A**). This outperformed models that used only covariates, or the 388 peaks identified through linear regression (P<0.001) (**Figure 2B**). Given the global hypoacetylation observed in this dataset, we reasoned that these 7 peaks may not be the only features capable of distinguishing opioid cases. To identify additional peak sets with similar predictive accuracy, we developed a stepwise strategy which iteratively removes peaks used by previous models and then identifies new models from the remaining peaks (**Figure 2C)**. In total, we identified 108 independent models ranging from 4 to 43 peaks with median AUCs >0.9 (**Supplemental Figure 2A,B, Table S3**). Altogether, these 108 models incorporated 3200 peaks. Using unbiased Uniform Manifold Approximation and Projection (UMAP) dimensionality reduction (McInnes et al., 2018), we found that these features could easily distinguish opioid cases from controls (**Figure 2D**) and that their separation was not observed when utilizing all ChIP-seq peaks (**Figure 2E**). Using DLPFC promoter-capture Hi-C data, we identified 830 gene targets of model peaks. While individual peaks were unique to each model, we found 167 (20%) target genes that were incorporated across multiple models. Genes found by three or more models include *DUSP4* and the potasium channel *KCNMA1* (**Figure 2F**). Using Functional Mapping and Annotation (FUMA) (Watanabe et al., 2017), gene targets of model peaks were enriched for genes broadly involved in synapse function and behavior, regulating GABAergic transmission, G protein signaling, and neuronal phosphodiesterases and sex steroid (incl. androgen) receptor pathways (**Figure 2G**). Collectively, these results demonstrate that while individual changes in regulatory elements are small, they are sufficient to distinguish opioid overdose death cases from controls. More importantly, the repeated appearance of synaptic gene targets in the 108 models, and the use of peaks beyond the 388 in these models, suggests the heterogeneity amongst individual cases is converging on shared underlying biology.

**Figure 2.**
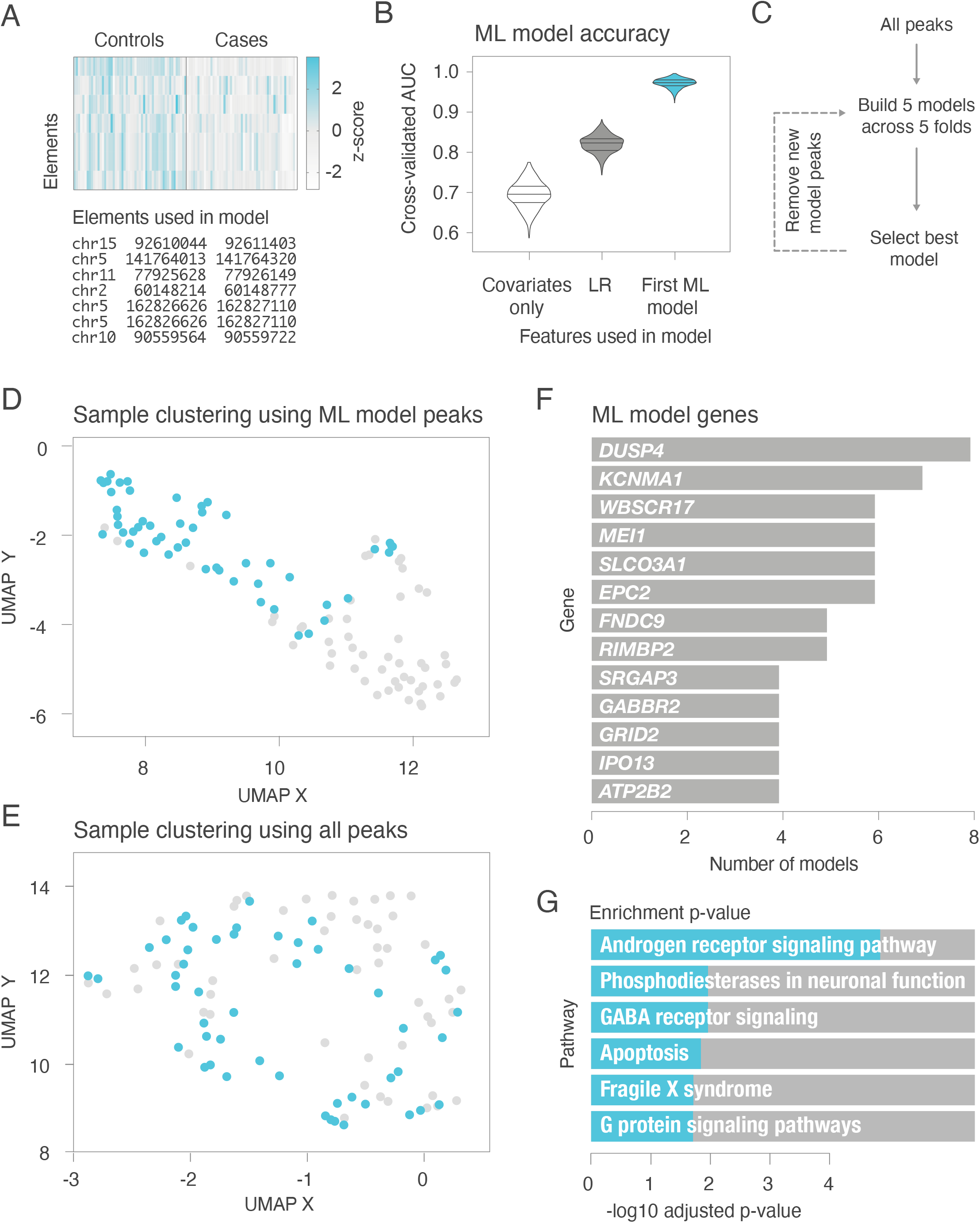
Machine learning reveals models and key features that distinguish opioid cases from controls. **(A)** Heatmap of the seven peaks that distinguish cases for controls in the first ML model. **(B)** Distribution of AUC derived from multiple 80/20 train/test splits (5-fold cross validation with 1,000 repetitions) shown for the top machine learning model (blue), models derived from the 388 linear regression peaks identified (grey) and models derived only using covariates (white). **(C)** Schematic representing iterative strategy to identify additional models with high predictive accuracy. **(D)** UMAP projection of opioid cases (blue) and control samples (grey) based on peaks utilized by 108 machine learning models and **(E)** based on all ChIP-seq peaks. **(F)** Gene targets that are shared across 4 or more ML models. **(G)** Pathways enriched across all ML model genes.

### Identification of heterogeneous case-specific alterations

In order to uncover the epigenetic heterogeneity of opioid dependence, we used an information-theoretic approach to identify changes that distinguish individual cases from the full set of controls (**Figure 3A, Methods**). A similar strategy has been used to pinpoint key regulatory differences among heterogeneous cancers (Akhtar-Zaidi et al., 2012)(Cohen et al., 2017)(Morrow et al., 2018). We refer to these individual-specific changes as Variable or Variant Enhancer Loci, or VELs. The vast majority of differentially acetylated regions identified through VEL-mapping were hypoacetylation events, consistent with the findings based on linear regression. Specifically, we identified 81,399 VELs with reduced H3K27ac, or “lost VELs”, in at least one opioid overdose death case relative to controls (FDR<5%), and 976 gained VELs (**Table S3**). Of the 388 hypoacetylated regions identified by linear regression (LR), all but two were detected via the VEL-mapping approach. On average the magnitude of the effect was far greater for H3K27ac differences found through VEL-mapping than linear regression (**Figure 3B**), suggesting that case-specific changes may be bigger contributors to phenotype than LR-defined events. VEL gene targets show changes in expression consistent with the accumulation of gains and losses, albeit asymmetrically, with gains showing a more immediate effect (**Figure 3C**). The number of VELs shared across cases is far greater for losses than gains (**Figure S3)**.

**Figure 3.**
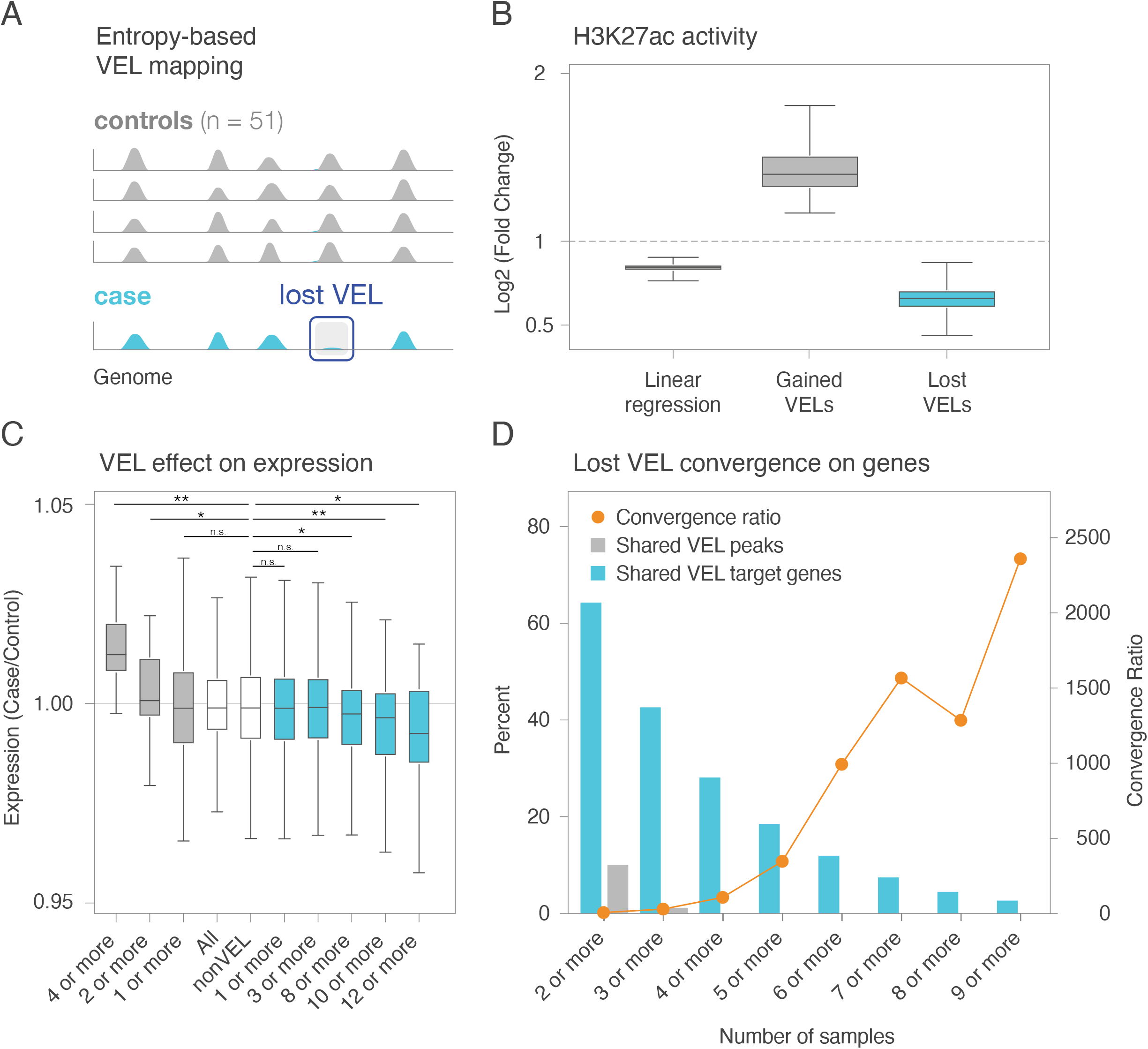
Identification of Variable Enhancer Loci (VELs) **(A)** Schematic overview of VEL identification. Each opioid case is compared to the full set of control samples to identify case-specific alterations in H3K27ac. **(B)** Log_2_ fold change (cases/controls) in ChIP-seq signal for lost VELs (blue), gained VELs (grey), and linear regression peaks (black). **(C)** Tukey boxplot of fold change in average gene expression for genes associated with VELs. Genes associated with gained VELs (grey) and lost VELs (blue) in multiple samples are shown as well as control gene lists shown in white. Outliers not shown. Kruskal-wallis ANOVA P<0.0001. Dunn’s post-hoc test is shown ^*^P<0.05 ^**^P<0.009. **(D)** Comparison of VELs and VEL gene targets across opioid cases. Percentage of VEL peaks shared across multiple cases (grey bar). Percentage of VEL gene targets shared across multiple samples (blue bar). Convergence ratio, i.e. the ratio of shared genes to shared VEL peaks (orange line).

Case-specific lost VELs show striking convergence at the gene level. Upon linking VELs to their putative gene targets using DLPFC promoter-capture Hi-C data, we find that while only 10% of lost VELs occur in two or more overdose cases, 60% of target genes were physically linked to VELs in two or more overdose cases (**Figure 3D**). As we look for VEL peaks and VEL target genes that are found in a larger number of cases, the percentage of genes and peaks decreases exponentially (**Figure 3D**). However, the ratio of VEL target genes to VEL peaks (convergence ratio) increases exponentially, indicating that heterogeneous case-specific losses accumulate at the gene level in a large number of cases. In other words, despite vast heterogeneity at the level of individual enhancers, the signal appears to “yield” at the gene level, becoming increasingly homogenous. We next set out to capture this homogeneity and resolve it into concrete genes at which these alterations converge through the development of a novel statistical framework which we refer to as “convergence analysis.”

### Regulatory losses converge on synaptic genes

Convergence analysis is designed to resolve genomically dispersed, low-frequency variation in either the DNA sequence or epigenomic signals into high-frequency convergent events. We use the term “plexus” to refer to the collection of regulatory elements that physically contact the gene promoter (Sallari et al., 2017)(Bailey et al., 2016). Plexi are derived from DLPFC promoter-capture Hi-C (Jung et al., 2019). Here, we use convergence analysis to identify genetic loci and target genes that have an overabundance of opioid case VELs. The strategy identifies single elements in a gene plexus altered across multiple cases, multiple elements altered within a single case, multiple elements altered across multiple cases, and intermediate patterns (**Figure 4A**). This approach takes into account the differing number of elements contained in each gene’s plexus and their activity to identify genes with the most significant accumulation of VELs.

**Figure 4.**
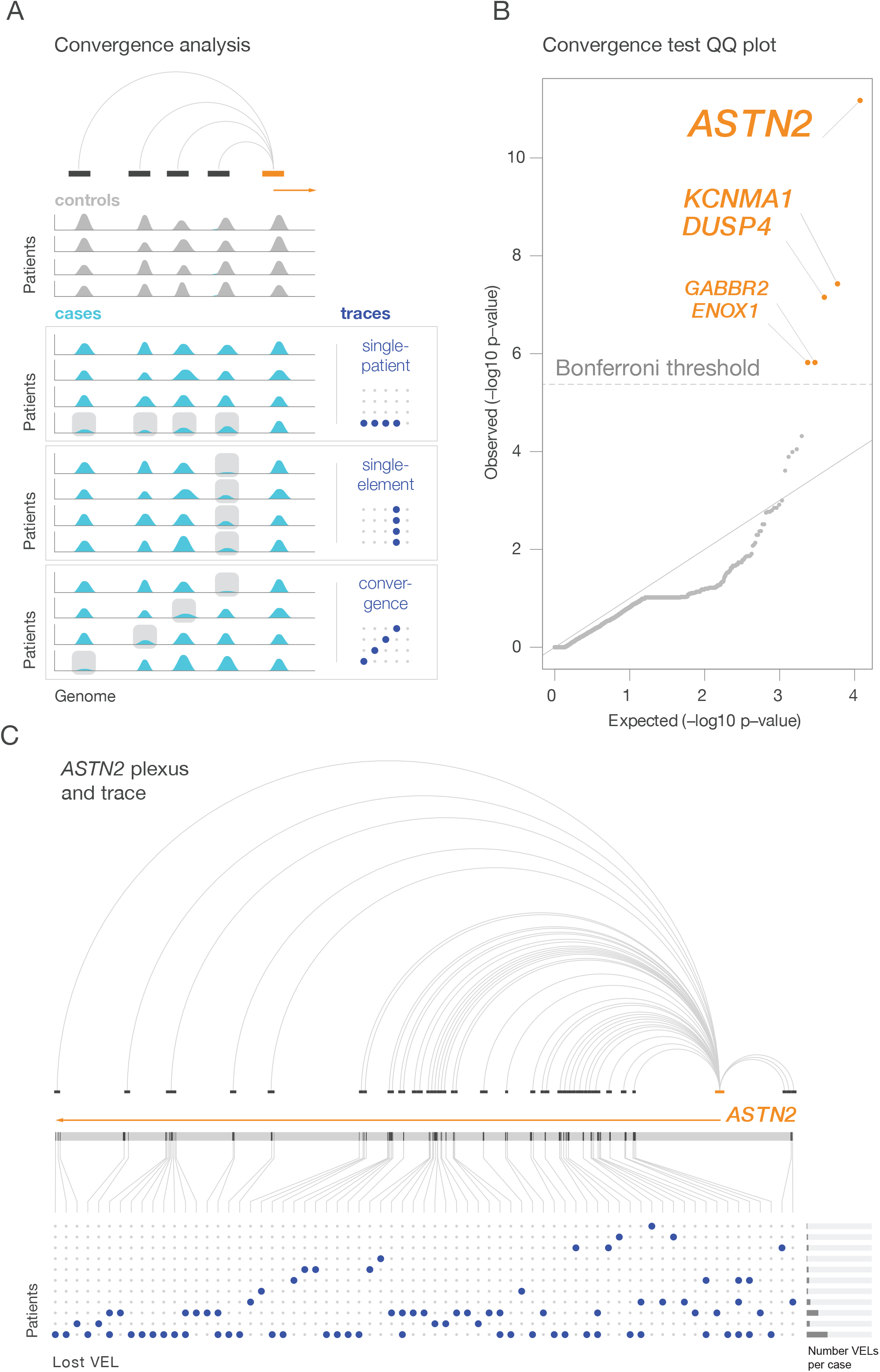
VEL convergence of shared target genes with plexus analysis. **(A)** Schematic representing plexus convergence strategy. (top) H3K27ac peaks (black) associated with common target promoter (orange) in promoter capture Hi-C (arcs). Example traces are shown for multiple VELs associated with the same target gene in one opioid case (single patient), multiple samples with the same VEL (single element), different VELs associated with the same target gene across cases (convergence). **(B)** Quantile-quantile plot of plexus analysis of VEL convergence. Five genome-wide significant genes are labelled (font proportional to p-value). **(C)** Plexus VEL trace for top gene *ASTN2* is shown with genomic location and Hi-C interaction indicated above. Bar graphs indicating the total number of VELs identified per case is shown (right).

Five convergent plexi are statistically significant for accumulation of lost VELs: *ASTN2, KCNMA1, DUSP4, GABBR2*, and *ENOX1* (*P*_*FDR*_: 1×10^−11.2^, 1×10^−7.4^, 1×10^−7.2^, 1×10^−5.8^, 1×10^−5.8^; **Figure 4B,C**). We do not identify significant plexi for the accumulation of gained VELs. The *ASTN2* plexus, the most significant result, is composed of 106 putative regulatory elements contained in loci that interact with the *ASTN2* promoter in DLPFC; 65.1% of its elements and 21.6% of the cases have at least one VEL (plexus and cohort convergence, respectively). *KCNMA1* shows the highest single-patient burden of losses: 36.3% (41 out of 113 elements). *DUSP4* shows the highest level of cohort convergence: 37.3% (19 of the 51 patients); however, in this plexus, the highest frequency of losses at any single element is 9.8% (5 of the 51 cases), highlighting the ability of the convergence framework to decode case heterogeneity. *GABBR2* shows the highest plexus convergence: 84.6% (88 out of 104 elements). The plexi of these five genes do not overlap, indicating that the significance of the results is mutually independent.

Here, we are able to distill 81,399 case-specific VEL alterations into five genes where global hypoacetylation is concentrated beyond what is expected by chance, showing clear erosion of regulatory activity across each gene’s plexus. Conventional strategies seek to identify robust changes that distinguish all cases from controls. The VEL convergence strategy offers a novel opportunity to take full advantage of the richness of information contained in the network of subtle and heterogenous regulatory changes.

### Convergent losses account for heritability of anxiety and overlap OUD risk genes

We hypothesized that direct ascertainment of converging heterogeneous effects of opioid exposure on the human brain’s epigenome will illuminate the genetic underpinnings of the traits involved in OUD. To evaluate this hypothesis, we compared the top 5 convergent plexi to the heritability of a panel of 27 traits (Finucane et al., 2015)(**Figure 5A,B, Table S6**). This included summary statistics from the MVP study of OUD with 10,544 cases and 72,163 controls with one replicable finding (*OPRM1*)(Zhou et al., 2020a). We did not observe heritability enrichment for OUD, potentially owing to low power of the OUD GWAS; alternatively, as *OPRM1* enhancers were not differentially acetylated, it is possible that opioid overdose death is epigenetically heritable at loci distinct from the genetically heritable loci. In evaluating other phenotypes, the top 5 gene’s plexi account for a significant proportion of heritability of generalized anxiety disorder (N=200K), number of sexual partners (N=370K) and educational attainment (N=126K)(**Figure 5C**). Given that regulatory regions active in the CNS have been previously shown to explain a substantial proportion of the heritability of neuropsychiatric disorders and behavioral phenotypes, we compared these results to control sets of cortical enhancers with similar H3K27ac patterns as the VELs. VELs associated with the top 5 plexus genes were substantially more enriched for heritability of these three phenotypes, with generalized anxiety disorder showing nearly 4-fold more heritability enrichment than activity-matched cortical enhancer regions (**Figure 5C**). These results demonstrate that previously identified genetic vulnerabilities to traits linked with opioid dependence co-localize with the convergent epigenetic changes we identified in our opioid overdose cohort. Notably, amongst all identified lost VELs, we observed heritability enrichment for additional traits including alcohol use disorder, drinks per week, smoking initiation, smoking cessation and measure of risk tolerance. However, in contrast to the top 5 plexi, heritability enrichment for VEL regions was equal to the enrichment found in other regulatory elements active in the neurons (**Supplemental Figure 4A,B)**.

**Figure 5.**
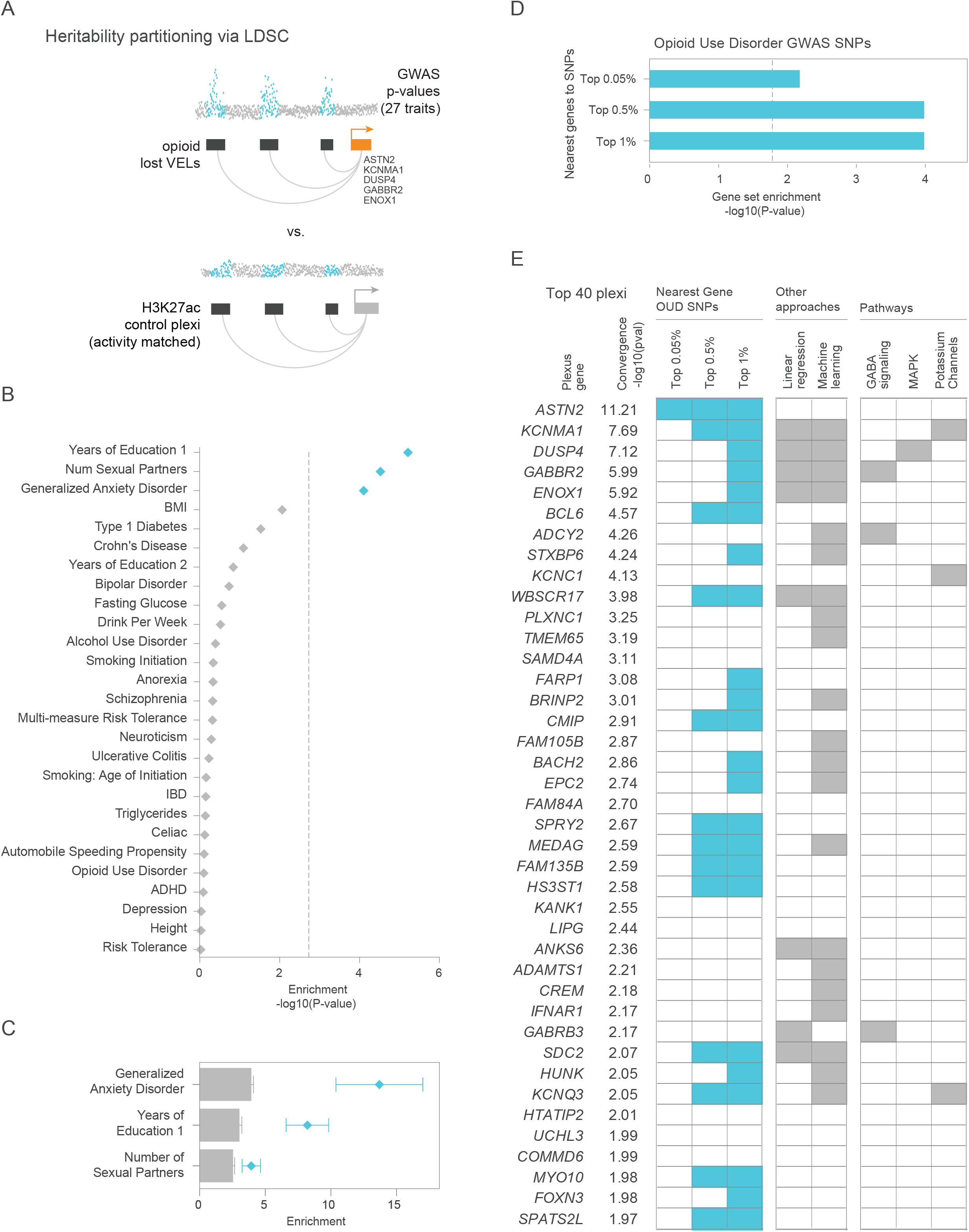
Convergent VEL genes are enriched for disease heritability. **(A)** Schematic representing analytical approach using LDSC to compare published GWAS results to top 5 plexus regions in (B) and to control regions with matched neuronal H3K27ac activity (C). **(B)** LDSC heritability enrichment (-log_10_ p-value) for VEL regions associated with top 5 plexus genes (Fig. 4B). Dotted line indicates Bonferroni multi-test correction threshold. Traits with significant enrichment are shown in blue. **(C)** Heritability enrichment (proportion of heritability / proportion of SNPs in peak set) with standard error is shown for VEL regions (blue dots) and 10 control sets of randomly selected neuronal peaks with activity profiles matched to VELs (grey bars). **(D)** GSEA analysis comparing genes ranked by plexus p-value to OUD candidate gene sets defined as genes nearest top 1%, 0.5% and 0.05% of SNPs identified by OUD GWAS (Zhou et al., 2020a). **(E)** Heatmap representing top 40 genes defined by plexus analysis (left). Genes nearest the top 1%, 0.5% and 0.05% of OUD GWAS SNPS are indicated (blue). P-value based on 10,000 permutations. Genes also identified by previous approaches including linear regression and machine learning models are highlighted in center. Genes linked to key pathways are indicated (right).

We expanded our analysis to the entire list of plexi ranked genes to further investigate overlap with OUD GWAS (Zhou et al., 2020a). We directly assessed whether loci with greater VEL convergence were enriched among the strongest OUD-associated SNPs as sorted by GWAS p-values. We identified genes nearest the top 1%, 0.5%, and 0.05% of OUD GWAS SNP associations. Using GSEA (Mootha et al., 2003; Subramanian et al., 2005), we found these genes to be significantly enriched amongst genes with the greatest VEL convergence (**Figure 5D**). Of the top 40 genes with the greatest VEL convergence, 23 (57.5%) were the nearest gene to a SNP in the top 1% of the OUD GWAS results (**Figure 5E**). Interestingly, of these 40 target genes, 8 were also identified through our earlier LR analysis and 21 through ML analysis. These 40 also include multiple genes from pathways enriched in previous analyses (**Figure 5E**). These results suggest that our multi-pronged approach may be effective at revealing loci that predispose to OUD and its related traits.

## Discussion

Neuropsychiatric traits like OUD and its most severe outcome, opioid overdose death, are complex and are best addressed by multiple complementary epigenomic analyses. A new approach like ours that embraces the deep heterogeneity of complex traits, is particularly valuable. We employ linear regression, ML, VEL mapping and convergence analysis to identify novel genes associated with opioid dependence. Our novel VEL-convergence approach recovers the rich, case-specific variation that would be discarded in a conventional study, either because their effects sizes would be too small in aggregate across subjects, scattered across the genome, or case-specific. The approach we use here leverages the topology of the genome to decode heterogeneous variation, resolving low-frequency patterns of variation across all regulatory elements connected to a gene into more detectable high-frequency alterations at the gene level. *DUSP4* provides a good example of the value of our ensemble approach: it is connected to an element identified by LR, and it shows a transcriptional change; however, it also harbors a preponderance of case-specific alterations, observed in 37.3% of the cases. Without the convergence approach, this profuse effect would have gone unnoticed. Most importantly, the convergence framework reveals additional genes with even more extensive burden of alterations that would have been completely missed, as they don’t fall near any of the 388 LR-identified elements.

We use three different strategies to characterize epigenetic events in opioid overdose cases. Across all three approaches, we identified *KCNMA1, DUSP4*, and *GABBR2* as critical target genes. Our analyses implicate MAPK signaling, GABA signaling and potassium channel genes. Each of these three pathways have been broadly linked to substance abuse and dependence. For example, KCNMA1 is a large conductance potassium channel previously linked to alcohol dependence (Bettinger and Davies, 2014) and considered a hub gene for alcohol-induced transcriptomic alterations in the prefrontal cortex (Wolen et al., 2012). *DUSP4* encodes a MAP kinase broadly relevant for synaptic plasticity and memory functions in the brain (Abdul Rahman et al., 2016). *GABBR2* encodes a metabotropic, G-protein coupled GABA receptor and potential OUD risk gene (Cui et al., 2012), with altered gene expression in the prefrontal cortex in an animal model (Ribeiro et al., 2012). Finally, common polymorphisms in ASTROTACTIN 2 (*ASTN2*), a membrane-bound regulator of the surface expression of many synaptic proteins (Behesti et al., 2018), exert a surprisingly strong effect on postoperative pain phenotypes, including patient-controlled dosing of opioid-based analgesics (Inoue et al., 2021).

The approach pioneered here is able to uncover some of the biology of OUD with as few as 51 cases and 51 controls - a sample size dwarfed by standard GWAS. Expanding to larger cohorts and other addiction-relevant brain regions beyond DLPFC may yield many more genes for further investigation as either biomarkers of opioid exposure, potential druggable targets, or causal genes that underlie risk of OUD, similarly to how one would functionally characterize GWAS hits. Coupling this approach to single cell strategies could further inform specific neuronal or non-neuronal cell subtypes that are especially affected by opioid abuse. Our approach does not replace GWAS and animal studies, as both are critical in advancing our understanding of complex traits, particularly for traits that disproportionately impact hard-to-reach populations. The hybrid experimental and computational strategy that we present here is designed to augment or even accelerate the mapping of complex genetic architectures.

What have we uncovered with this new methodology, and what is the relevance to the current opioid crisis? We believe our study is capturing shards of the genetic and environmental factors perpetuating the opioid epidemic. In fact, our top 5 plexus-genes are enriched among the top 1% of OUD GWAS. While we did not observe global enrichment for OUD heritability, this is not unexpected given the incompleteness of the OUD genetic architecture, as the largest OUD GWAS captures <12% of the expected heritability. Alternatively, the most severe outcome of OUD, opioid overdose death, may be driven by genetic and/or environmental factors that are either entirely distinct from OUD, or are comprised of a particular subset of OUD risk factors. This further highlights the need for approaches which simultaneously captures the genetics and epigenetic influences of the disorder.

The three traits uncovered in the top 5 gene plexi - anxiety, years of education and number of sexual partners - have been associated with early childhood adversity (Hillis et al., 2001)(Montez and Hayward, 2014)(Murthy and Gould, 2020). These phenotypes as well as early childhood adversity are major risk factors for OUD (Levis et al., 2019)(Enoch, 2011)(Widom et al., 2006). This observation points to a potential shared pleiotropic risk that may be revealed as the risk factors for all of these related traits are uncovered. Early life adversity (ELA) and resource scarcity has been shown to have lasting impacts on the epigenome in both rodent and human studies (Vaiserman and Koliada, 2017)(Barnett Burns et al., 2018), and thus has the potential to play a causal role in some of the epigenetic changes we identified. ELA is an increasingly recognized contributor to many public health crises (Leza et al., 2021). We expect that genetic and environmental factors, such as ELA and drug exposure, may cooperatively contribute to the epigenetic changes we have identified. Risk factors in adulthood for opioid overdose death, such as lack of social support, exposure to opioids, and lacking economic opportunity have been rising steadily over the past few decades, driven by shifting socioeconomic factors and changes in opioid availability (Dasgupta et al., 2018)(Altekruse et al., 2020). Angus Deaton and colleagues called these changes “the ongoing erosion of working-class life for those born after 1950” (Case et al., 2020). In their landmark study, they describe a generational increase in self-reported pain in the United States, which they tie to the opioid epidemic. Intriguingly, two variants in our top result, *ASTN2*, are associated with sensitivity to opioids in a study of pain-related phenotypes of fentanyl use after surgery (Inoue et al., 2021), suggesting we may also be identifying variation in pain response. Furthermore, genetic and environmental risk factors may influence the overlapping sets of target genes through distinct mechanisms. Epigenomic studies like ours are uniquely poised to identify the loci most critically affected by this milieu of genetic and environmental influences.

Our study identifies novel elements of the OUD exposure and risk architecture, as well as phenotypes related to early life adversity, expanding the field’s current limited view of the underlying influences that contribute to the opioid epidemic. Altogether our results indicate that we are seeing just the tip of the iceberg of the intertwined causes, consequences of opioid exposure in the brain, and related traits that ultimately lead to opioid related fatalities. The epigenome is a palimpsest where inherited epigenetic and genetic patterns are repeatedly overwritten by environmental exposures throughout our lives. The ensemble of methods we present here allow us to reveal previously-hidden genes. Further studies are needed to distinguish the nature of the factors leading to their dysregulation. Expanding both epigenetic and genetic studies of OUD and related phenotypes will enable us to further elucidate how diverse contributors manifest in the brain and drive this devastating epidemic.

## Supplementary Figures

**Supplemental Figure 1:**
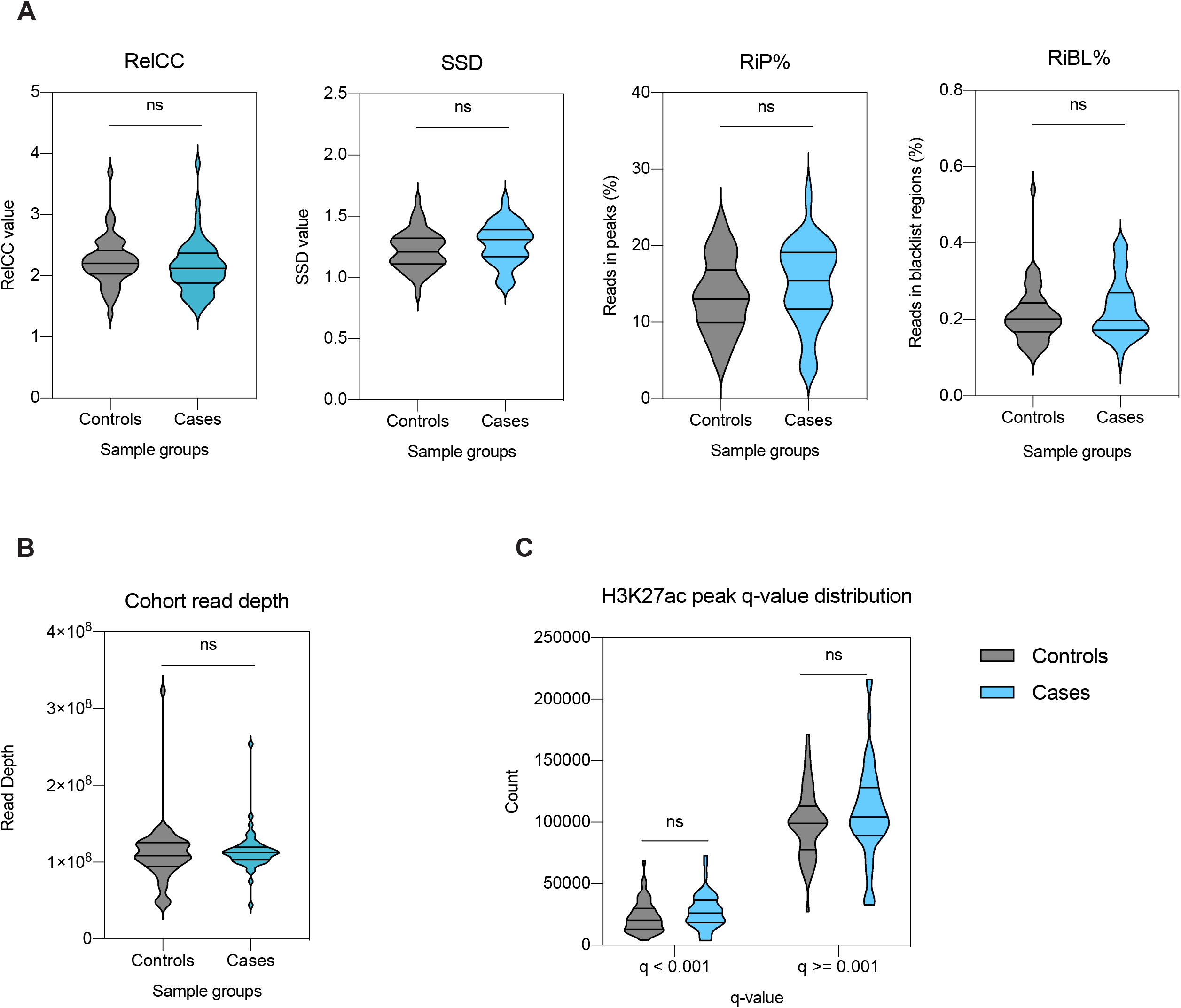
ChIP-Seq quality metrics. **(A)** ChIPQC metrics for the cases and controls show high similarity and quality between sample groups. Samples with RelativeCC (RelCC) > 1 and reads in peak percentage (RiP%) > 5% indicate good enrichment, while similar SSD values between sample groups indicate consistent quality. All samples had a small percentage of reads in blacklisted regions (RiBL%) < 0.54%. **(B)** Sample read depth between cases and controls. **(C)** Bonferroni multi-test-corrected p-values (q-values) show similar distributions of significant (q-value < 0.001) and less significant (q-value >= 0.001) H3K27ac peaks across sample groups.

**Supplemental Figure 2:**
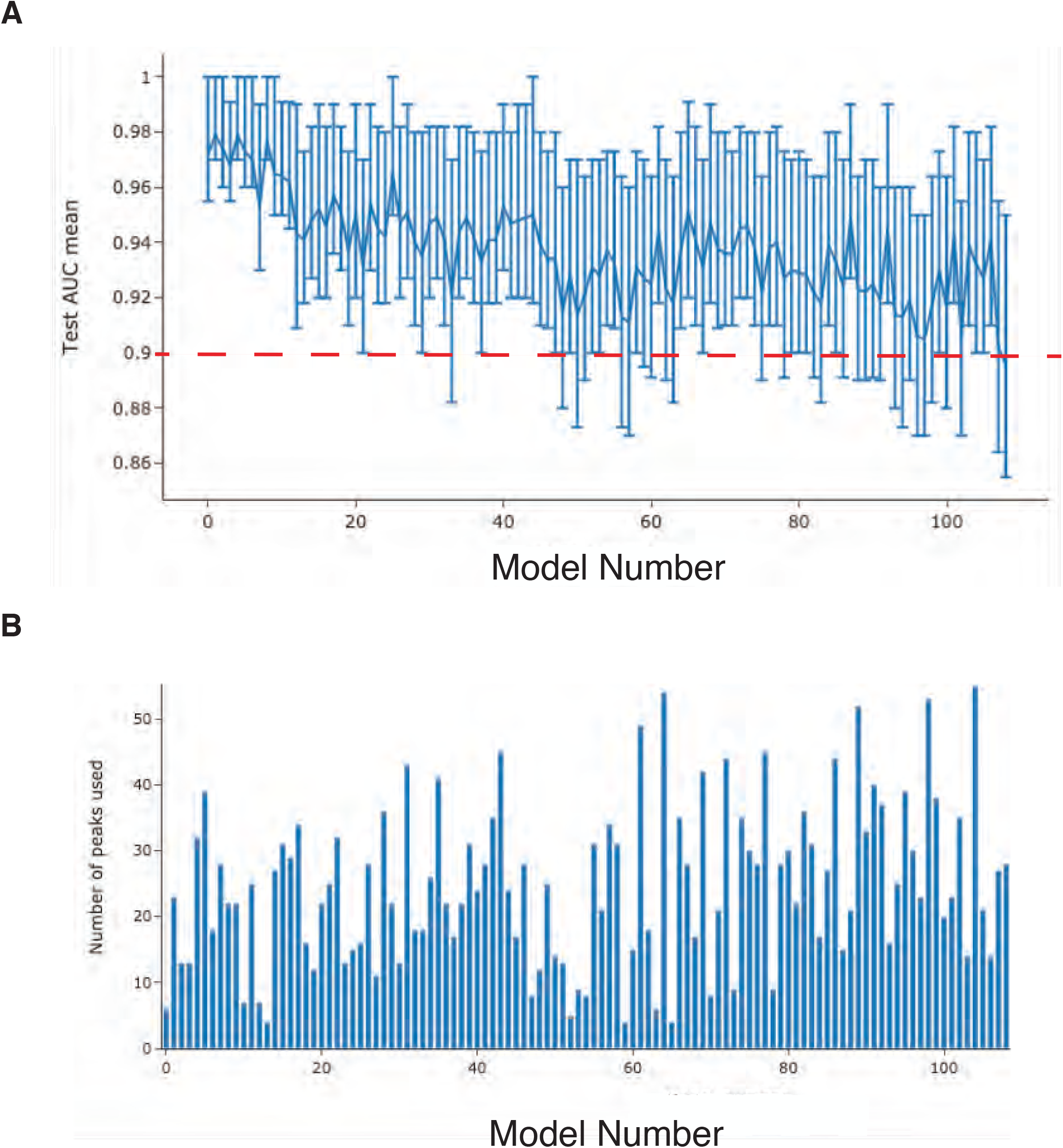
108 machine learning models description. **(A)** AUC distribution for 108 ML models generated. Shown are median in interquartile range for each model in order. **(B)** Number of peaks (features) utilized by each model.

**Supplemental Figure 3:**
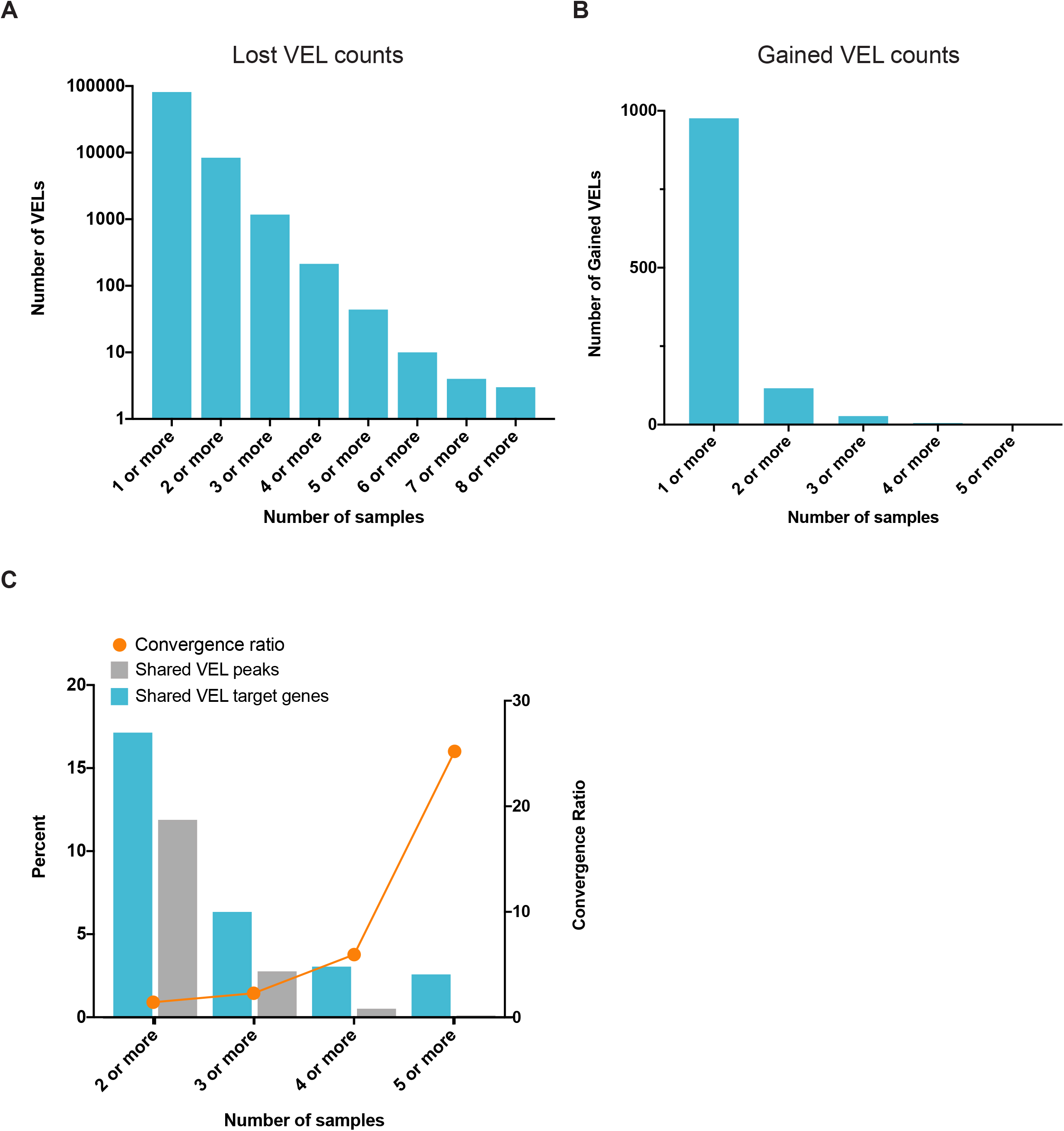
VEL counts and Gained VEL convergence. Number of VELs and shared VELs identified for lost VELs **(A)** and gained VELs **(B). (C)** Comparison of VELs and VEL gene targets across opioid cases. Percentage of VEL peaks shared across multiple cases (grey bar). Percentage of VEL gene targets shared across multiple samples (blue bar). Convergence ratio, i.e. the ratio of shared genes to shared VEL peaks (orange line).

**Supplemental Figure 4:**
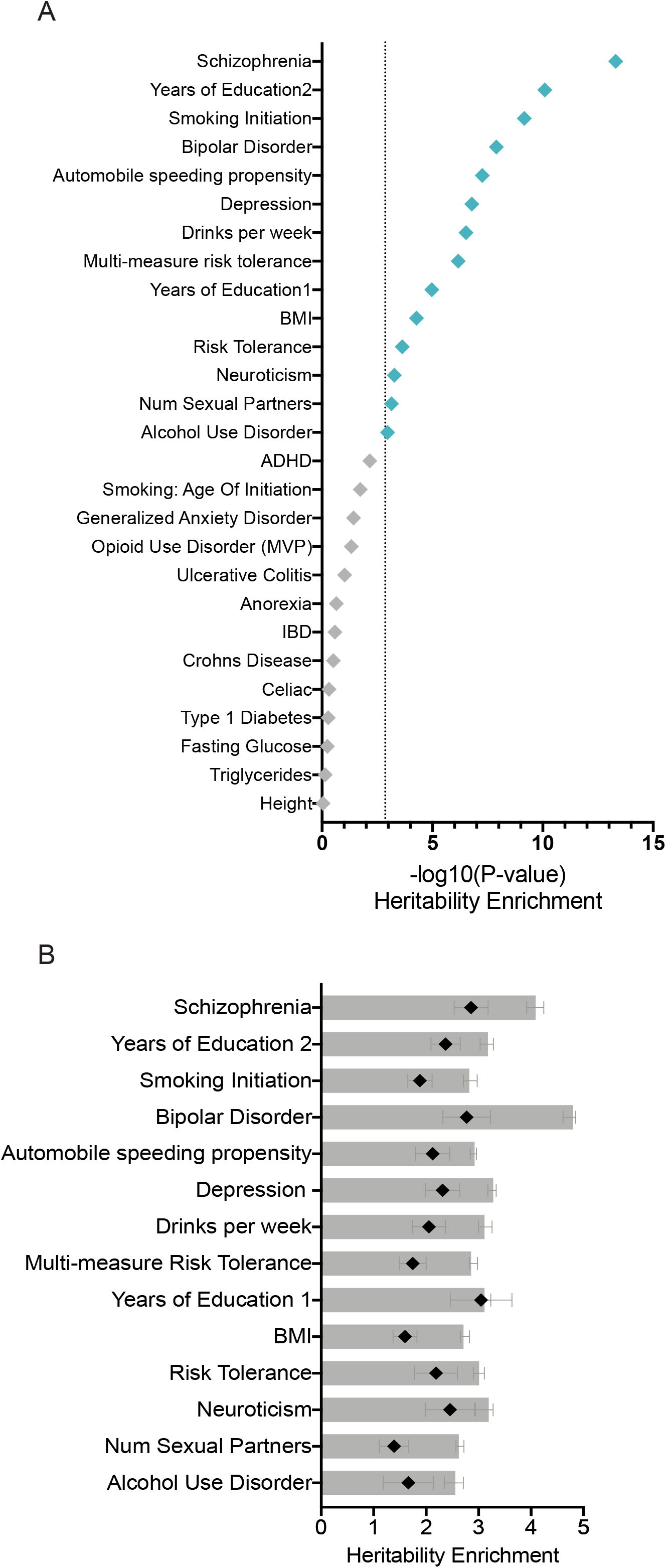
Heritability enrichment in all lost VELs. **(A)** LDSC heritability enrichment (-log_10_ p-value) for all lost VEL regions. Dotted line indicates Bonferroni multi-test correction threshold. Traits with significant enrichment are shown in blue. **(C)** Heritability enrichment (proportion of heritability / proportion of SNPs in peak set) with standard error is shown for VEL regions (black dots) and 10 control sets of randomly selected neuronal peaks with activity profiles matched to VELs (grey bars).

**Supplemental Table 1:** Panel description

**Supplemental Table 2:** Differentially acetylated peaks

**Supplemental Table 3:** Features utilized by machine learning models

**Supplemental Table 4:** Variable Enhancer Loci

**Supplemental Table 5:** Plexus results

**Supplemental Table 6:** Summary of GWAS studies utilized in Fig. 5.

## METHODS

### Patient cohort

Cases (N= 51) were selected from an opportunistic sample of opioid-related deaths and unaffected controls that came to autopsy. The use of de-identified cadaver specimens is not defined as human subjects research and exempt from regulation 45 CFR pt 46 (NIH SF424 guide Part II: Human Subjects). The cause of death was determined by forensic pathologists following medico-legal investigations evaluating the circumstances of death including medical records, police reports, autopsy findings and forensic toxicology analysis. Common drugs of abuse and alcohol and positive urine screens were confirmed by quantitative analysis of blood and brain. The detection of 6-acetyl morphine (6-AM) was taken as definitive evidence of acute heroin exposure. Retrospective chart reviews were conducted to confirm history of opioid abuse, methadone or addiction treatment, drug related arrests or drug paraphernalia found at the scene. Supplemental brain toxicology was done on select cases for comparison to blood levels at the time of death. Case inclusion in the opioid group was based on a documented history of opioid abuse, toxicology report positive for opioids, and forensic determination of opioids as cause of death. Drug-free control subjects (N=51) were age-matched from sudden accidental (motor vehicle accidents or trauma) or cardiac deaths with negative urine screens for all common drugs and no history of licit or illicit drug use reported by next-of-kin informants prior to death. Exclusion criteria for either group included death by suicide, a known history of psychiatric disorder, or debilitating chronic illness. The subject groups were matched as closely as possible for age, gender, postmortem interval (PMI), and agonal factors.

### ChIP-Seq data processing

Cutadapt v1.9.1 (Martin, 2011) was used to remove paired-end adapter sequences and discard reads with a length less than 20bp. All fastqs were aligned to the hg19 genome assembly (retrieved from hgdownload.cse.ucsc.edu/goldenPath/hg19/chromosomes) using BWA-MEM v0.7.17-r1188 (Li, 2013) with default parameters in paired-end mode. Output SAM files were converted to binary (BAM) format, sorted, indexed, and PCR duplicates were removed using SAMtools v1.10 (Li et al., 2009). Peaks were detected with MACS v2.1.2 (Zhang et al., 2008) with the --broad flag set. DeepTools v3.2.0 (Ramírez et al., 2016) was used to generate RPGC-normalized bigWig tracks with 50 bp bin sizes of the final sample BAM files and visualized on the Integrative Genomics Viewer (Thorvaldsdóttir et al., 2013) in order to eliminate samples with pronounced track irregularities or low signal-to-noise ratio. The Bioconductor (Gentleman et al., 2004) package ChIPQC (Carroll et al., 2014) was used to generate quality metrics for each processed library – those libraries which contained very few mapped reads, a RelativeCC enrichment score (RelCC) < 1, percentage of reads in peak regions (RiP%) < 2%, or number of detected peaks < 10,000 were excluded from additional analysis. Peak lists were filtered to remove all peaks overlapping ENCODE blacklisted regions (https://sites.google.com/site/anshulkundaje/projects/blacklists).

### Normalization

We use the Inherent Normalization algorithm as part of the Axiotl Convergence Platform to process and normalize H3K27ac BigWig tracks. This normalization approach is designed specifically for epigenomes and uses a robust and reproducible reference found within the sample itself. In order to find this “inherent unit” we first focus on the most stable epigenomic elements in the genome: promoters. We use the set of 81,232 reference promoters from the Roadmap project [Roadmap2015]. This set of promoters contains a subset of constitutive elements (23,374), active in virtually all cell types, and a subset of highly cell-type-specific elements (16,807), active in only one reference cell type. We refer to the first list as the “on” promoters and to the second as the “off” promoters. By using 111 reference epigenomes, the group of “off” promoters will overwhelmingly be inactive in any cell type. Given the large number of elements in each list, we can compute very precise averages. The distance between the averages of the “on” and “off” collections constitutes our “inherent unit,” hence the name of the algorithm. The values of the “on” and “off” averages in each sample are then used to align the data, such that all the “off” averages across samples are at zero and all “on” averages are at one. Data is log2 transformed so as to make the data normally distributed. The approach is described in more detail in Chakraborty et al. (In preparation), where the authors show consistent reduction of noise when reprocessing the ENCODE 3 reference epigenomes. Noise reduction translates to greater statistical power to discover new results in the data. The approach also makes the data biologically interpretable and universally meaningful, with a value of one representing the standard activity of a constitutive promoter.

### RNA-seq data processing and quality control

Postmortem, dorsolateral prefrontal cortex (Brodmann area 9) were dissected from cryopreserved coronal slices of brain to obtain bulk tissue RNA samples. 150 samples were prepared for sequencing using Takara Bio SMARTer Stranded RNA-seq Kit and 150 bp paired-end sequencing was performed on an Illumina. *Trimmomatic v0*.*39 (Bolger et al., 2014)* was used for read trimming and quality filtering with the following settings: *HEADCROP length*=3, *ILLUMINACLIP seedMismatches*=2, *ILLUMINACLIP palindromeClipThreshold*=30, *ILLUMINACLIP simpleClipThreshold*=10, *ILLUMINACLIP minAdapterLength*=4, *ILLUMINACLIP keepBothReads*=true, *LEADING quality*=3, *TRAILING quality*=3, *SLIDINGWINDOW windowSize*=4, *SLIDINGWINDOW requiredQuality*=10, *MINLEN length*=75, *AVGQUAL quality*=20. Read pairs with both reads passing quality filtering were mapped using *Salmon v1*.*1*.*0 (Patro et al., 2017)* in selective alignment mode (Srivastava et al., 2020) with an index comprised of a GENCODE v30 transcriptome (comprehensive gene annotation) and the full GRCh38 primary assembly as a decoy. The options *--gcBias* and *--seqBias* were enabled to account for potential transcript quantification biases (Love et al., 2016). Salmon transcript quantifications were aggregated to the gene level using *tximport* v1.12.3 (Soneson et al., 2015).

Samples were excluded based on the following criteria: >40% of reads removed by Trimmomatic, mean read GC content <40%, effective sequencing depth <10 million read pairs, transcriptomic mapping rate <0.3, RIN score <5, mitochondrial mapping rate >0.5, gene expression principal component analysis outliers (along principal components 1 and 2), age outlier (age at death >60 years), and self-reported race not African-American or White. Several RNA samples were derived from the same individuals (i.e., replicates). For each of these replicates sets, the sample with the highest effective sequencing depth was retained among the samples that passed quality control. Following all sample exclusions, 24 opioid intoxication cases and 27 controls were used for subsequent analysis.

### Differential gene expression analysis

Genes were tested for differential expression by opioid overdose death case/control status using *DESeq2 v1*.*26*.*0 (Love et al., 2014)* to fit a generalized linear model with normalized gene counts as the outcome. Opioid intoxication status, ancestry, sex, age at death, PMI, RIN, RNA-seq sequencing batch, and 11 surrogate variables (SVs) were included in the model as explanatory variables. The 11 SVs were estimated using *SVA v3*.*36*.*0 (Leek et al., 2012)* with variance stabilized gene counts and the aforementioned explanatory variables as null model variables. To reduce the multiple testing burden, lowly expressed genes, defined as having <10 counts in 40% of samples (i.e., the approximate proportion of cases), were excluded, leaving 20,855 genes in our analysis.

### Variable Enhancer Loci (VEL) mapping approach

VELs were identified as previously described (Akhtar-Zaidi et al., 2012). Briefly, each opioid case was compared independently to the full panel of controls. Shannon entropy scoring was performed on inherent normalized H3K27ac ChIP-seq data (described above) to quantify peak specificity for each opioid case. Case/control status was then randomly permuted and shannon entropy scoring was repeated. These permutations were performed at least 70,000 times to generate a unique null expectation of peak specificity scores for each individual peak region. These permutations reveal the expected distribution of specificity scores due to inter-sample variation at each peak region. The true value was compared to these distributions for each peak to define the p-value. As each peak was tested twice to identify cases with significantly higher signal than controls (gained VELs) and significantly lower (lost) VELs, p-values were first multiplied by 2. We then applied Benjamini-Hochberg FDR correction to identify lost and gained VELs at an FDR 0.05%.

### Machine learning models

We utilized inherent normalized (see above) data for all 652,055 H3K27ac peaks, as well as a list of covariates as input features. The covariates include: sample descriptors such as, sex, age, ancestry and PMI; and ChIP-seq data descriptors such as number of peaks per ChIP, number sequencing reads generated, number of aligned reads after duplicate removal, alignment rate, fraction reads in peaks, and batch number. We used XGBoost’s (Chen and Guestrin, 2016) default hyper-parameters to generate models to predict case control status of ChIP samples and only varied the input feature list in order to compare the models.

#### Step-wise strategy for peak set removal

We utilized a step-wise strategy to identify distinct models capable of distinguishing cases and controls at high accuracy. The first model receives all input features (see definition above). For each subsequent model, the features utilized by the previous models are removed from the input feature list. We repeated this strategy until we identified a model where the AUC median (described below) fell below 0.9, resulting in a total of 108 models. To find peak sets that are predictive and robust to the choice of train/test set, we utilized two stage assessment of new models: 1) peak set discovery and 2) peak set evaluation.

#### Peak set discovery

We split the data into five folds, with 4 folds of 10 samples and 1 fold of 11 samples. We train 5 models, each using a different combination of four folds as training set (80% of samples) and the remaining fold as test set (20% of samples). By training 5 models, each fold is used as test set exactly once. Early stopping was used during training to obtain the minimal number of features that achieves good performance. After training, we examined each model to extract information about which features the model deemed most relevant to the classification task. We used the SHAP package to calculate SHAP values for each feature per sample (Lundberg et al., 2020). In order to find features that are important in distinguishing cases from controls, we used the mean of absolute SHAP values as the overall importance of each feature across all samples (a non-negative value). We define the “peak set” of a model as peaks that had non-zero overall importance.

#### Peak set evaluation

The predictive performance of a peak set in stage 1 might be specific to how the samples are split into folds. If there is heterogeneity in the data, the specific samples in the test fold can influence model performance. Thus, to obtain unbiased metric for model performance, we evaluated the peak sets on multiple different 5-fold splits. For each of the 5 peak set discovered in step 1, we trained 1000 5-fold cross-validated models (5000 models in total per peak set). To select the final peak set, we examine the AUC distribution across the aforementioned models. The chosen peak set for downstream analyses is the one that ranks highest when sorted by decreasing AUC mean and AUC minimum.

### Convergence analysis

Interactions between gene promoters and non-promoter loci are intersected with merged, non-overlapping elements called from the 51 case and 51 control H3K27ac tracks (652,055 elements).

Convergence analysis is run on two binary matrices of gained and lost variant H3K27ac elements (VEL) identified on 51 opioid-overdose individuals using our entropy-based approach using the Axiotl Convergence Platform (**convergence**.**axiotl**.**com**). The analysis consists of a permutation test in which a convergence statistic in the observed plexus is compared to a null distribution obtained from a collection of null plexi sampled from the binary matrix so as to match the activity and connectivity properties of the original plexus. Heterogeneity in global patient activity is also preserved. The algorithm is described in detail in Sallari et al. (manuscript in preparation).

### LD score regression

LDSR was used to estimate heritability enrichment across significant plexi. Annotated SNPs lists were generated for all SNPs within VEL regions (+/-5kb from VEL peak centers) associated with the top 5 significant plexi. European LD scores and weights were downloaded from the LDSC (Bulik-Sullivan et al., 2015). Summary statistics for 27 traits were obtained from previously published studies listed in Supplemental Table S6. LD scores were calculated as recommended https://github.com/bulik/ldsc/wiki/ using HapMap3 SNPs and 1000 genomes phase 3 reference files (1000 Genomes Project Consortium et al., 2015). Heritability enrichment calculations were performed using the recommended ‘baseline model’, composed of multiple functional categories including conserved regions and coding regions. The significance threshold was defined through Bonferroni multi-test correction for 27 traits tested.

## Notes

### Competing Interest Statement

The authors have declared no competing interest.

## References

1000 Genomes Project Consortium, Auton, A., Brooks, L.D., Durbin, R.M., Garrison, E.P., Kang, H.M., Korbel, J.O., Marchini, J.L., McCarthy, S., McVean, G.A., et al. (2015). A global reference for human genetic variation. Nature 526, 68–74.

Abdul Rahman, N.Z., Greenwood, S.M., Brett, R.R., Tossell, K., Ungless, M.A., Plevin, R., and Bushell, T.J. (2016). Mitogen-Activated Protein Kinase Phosphatase-2 Deletion Impairs Synaptic Plasticity and Hippocampal-Dependent Memory. J. Neurosci. 36, 2348–2354.

Akhtar-Zaidi, B., Cowper-Sal-lari, R., Corradin, O., Saiakhova, A., Bartels, C.F., Balasubramanian, D., Myeroff, L., Lutterbaugh, J., Jarrar, A., Kalady, M.F., et al. (2012). Epigenomic enhancer profiling defines a signature of colon cancer. Science 336, 736–739.

Al-Hasani, R., and Bruchas, M.R. (2011). Molecular mechanisms of opioid receptor-dependent signaling and behavior. Anesthesiology 115, 1363–1381.

Allen, H.L., Estrada, K., Lettre, G., Berndt, S.I., Weedon, M.N., Rivadeneira, F., Willer, C.J., Jackson, A.U., Vedantam, S., Raychaudhuri, S., et al. (2010). Hundreds of variants clustered in genomic loci and biological pathways affect human height. Nature 467, 832–838.

Altekruse, S.F., Cosgrove, C.M., Altekruse, W.C., Jenkins, R.A., and Blanco, C. (2020). Socioeconomic risk factors for fatal opioid overdoses in the United States: Findings from the Mortality Disparities in American Communities Study (MDAC). PLoS One 15, e0227966.

Bailey, S.D., Desai, K., Kron, K.J., Mazrooei, P., Sinnott-Armstrong, N.A., Treloar, A.E., Dowar, M., Thu, K.L., Cescon, D.W., Silvester, J., et al. (2016). Noncoding somatic and inherited single-nucleotide variants converge to promote ESR1 expression in breast cancer. Nat. Genet. 48, 1260–1266.

Barnett Burns, S., Almeida, D., and Turecki, G. (2018). The Epigenetics of Early Life Adversity: Current Limitations and Possible Solutions. In Progress in Molecular Biology and Translational Science, D.R. Grayson, ed. (Academic Press), pp. 343–425.

Behesti, H., Fore, T.R., Wu, P., Horn, Z., Leppert, M., Hull, C., and Hatten, M.E. (2018). ASTN2 modulates synaptic strength by trafficking and degradation of surface proteins. Proc. Natl. Acad. Sci. U. S. A. 115, E9717–E9726.

Bettinger, J.C., and Davies, A.G. (2014). The role of the BK channel in ethanol response behaviors: evidence from model organism and human studies. Front. Physiol. 5, 346.

Blackwood, C.A., McCoy, M.T., Ladenheim, B., and Cadet, J.L. (2021). Oxycodone self-administration activates the mitogen-activated protein kinase/ mitogen-and stress-activated protein kinase (MAPK-MSK) signaling pathway in the rat dorsal striatum. Sci. Rep. 11, 2567.

Bolger, A.M., Lohse, M., and Usadel, B. (2014). Trimmomatic: a flexible trimmer for Illumina sequence data. Bioinformatics 30, 2114–2120.

Boraska, V., Franklin, C.S., Floyd, J.A.B., Thornton, L.M., Huckins, L.M., Southam, L., Rayner, N.W., Tachmazidou, I., Klump, K.L., Treasure, J., et al. (2014). A genome-wide association study of anorexia nervosa. Mol. Psychiatry 19, 1085–1094.

Bradfield, J.P., Qu, H.-Q., Wang, K., Zhang, H., Sleiman, P.M., Kim, C.E., Mentch, F.D., Qiu, H., Glessner, J.T., Thomas, K.A., et al. (2011). A Genome-Wide Meta-Analysis of Six Type 1 Diabetes Cohorts Identifies Multiple Associated Loci. PLoS Genet. 7, e1002293.

Browne, C.J., Godino, A., Salery, M., and Nestler, E.J. (2020). Epigenetic Mechanisms of Opioid Addiction. Biol. Psychiatry 87, 22–33.

Brynildsen, J.K., Mace, K.D., Cornblath, E.J., Weidler, C., Pasqualetti, F., Bassett, D.S., and Blendy, J.A. (2020). Gene coexpression patterns predict opiate-induced brain-state transitions. Proc. Natl. Acad. Sci. U. S. A. 117, 19556–19565.

Bulik-Sullivan, B.K., Loh, P.-R., Finucane, H.K., Ripke, S., Yang, J., Schizophrenia Working Group of the Psychiatric Genomics Consortium, Patterson, N., Daly, M.J., Price, A.L., and Neale, B.M. (2015). LD Score regression distinguishes confounding from polygenicity in genome-wide association studies. Nat. Genet. 47, 291–295.

Carroll, T.S., Liang, Z., Salama, R., Stark, R., and de Santiago, I. (2014). Impact of artifact removal on ChIP quality metrics in ChIP-seq and ChIP-exo data. Front. Genet. 5, 75.

Case, A., Deaton, A., and Stone, A.A. (2020). Decoding the mystery of American pain reveals a warning for the future. Proc. Natl. Acad. Sci. U. S. A. 117, 24785–24789.

Center for Disease Control (2020). Increase in fatal drug overdoses across the United States driven by synthetic opioids before and during the COVID-19 pandemic. Publication Number CDCHAN-00438. December Available at https://emergency.Cdc.gov/han/2020/han00438.Asp (Accessed February 14, 2021).

Chen, T., and Guestrin, C. (2016). XGBoost: A Scalable Tree Boosting System. In Proceedings of the 22nd ACM SIGKDD International Conference on Knowledge Discovery and Data Mining, (New York, NY, USA: Association for Computing Machinery), pp. 785–794.

Chen, E.Y., Tan, C.M., Kou, Y., Duan, Q., Wang, Z., Meirelles, G.V., Clark, N.R., and Ma’ayan, A. (2013). Enrichr: interactive and collaborative HTML5 gene list enrichment analysis tool. BMC Bioinformatics 14, 128.

Cohen, A.J., Saiakhova, A., Corradin, O., Luppino, J.M., Lovrenert, K., Bartels, C.F., Morrow, J.J., Mack, S.C., Dhillon, G., Beard, L., et al. (2017). Hotspots of aberrant enhancer activity punctuate the colorectal cancer epigenome. Nat. Commun. 8, 1–13.

Cui, W.-Y., Seneviratne, C., Gu, J., and Li, M.D. (2012). Genetics of GABAergic signaling in nicotine and alcohol dependence. Hum. Genet. 131, 843–855.

Dasgupta, N., Beletsky, L., and Ciccarone, D. (2018). Opioid Crisis: No Easy Fix to Its Social and Economic Determinants. Am. J. Public Health 108, 182–186.

Demontis, D., Walters, R.K., Martin, J., Mattheisen, M., Als, T.D., Agerbo, E., Baldursson, G., Belliveau, R., Bybjerg-Grauholm, J., Bækvad-Hansen, M., et al. (2019). Discovery of the first genome-wide significant risk loci for attention deficit/hyperactivity disorder. Nat. Genet. 51, 63–75.

Dubois, P.C.A., Trynka, G., Franke, L., Hunt, K.A., Romanos, J., Curtotti, A., Zhernakova, A., Heap, G.A.R., Adány, R., Aromaa, A., et al. (2010). Multiple common variants for celiac disease influencing immune gene expression. Nat. Genet. 42, 295–302.

Enoch, M.-A. (2011). The role of early life stress as a predictor for alcohol and drug dependence. Psychopharmacology 214, 17–31.

Finucane, H.K., Bulik-Sullivan, B., Gusev, A., Trynka, G., Reshef, Y., Loh, P.-R., Anttila, V., Xu, H., Zang, C., Farh, K., et al. (2015). Partitioning heritability by functional annotation using genome-wide association summary statistics. Nat. Genet. 47, 1228–1235.

Gentleman, R.C., Carey, V.J., Bates, D.M., Bolstad, B., Dettling, M., Dudoit, S., Ellis, B., Gautier, L., Ge, Y., Gentry, J., et al. (2004). Bioconductor: open software development for computational biology and bioinformatics. Genome Biol. 5, R80.

Hancock, D.B., Markunas, C.A., Bierut, L.J., and Johnson, E.O. (2018). Human Genetics of Addiction: New Insights and Future Directions. Curr. Psychiatry Rep. 20, 8.

Hillis, S.D., Anda, R.F., Felitti, V.J., and Marchbanks, P.A. (2001). Adverse childhood experiences and sexual risk behaviors in women: a retrospective cohort study. Fam. Plann. Perspect. 33, 206–211.

Howard, D.M., Adams, M.J., Clarke, T.-K., Hafferty, J.D., Gibson, J., Shirali, M., Coleman, J.R.I., Hagenaars, S.P., Ward, J., Wigmore, E.M., et al. (2019). Genome-wide meta-analysis of depression identifies 102 independent variants and highlights the importance of the prefrontal brain regions. Nat. Neurosci. 22, 343–352.

Inoue, R., Nishizawa, D., Hasegawa, J., Nakayama, K., Fukuda, K.-I., Ichinohe, T., Mieda, T., Tsujita, M., Nakagawa, H., Kitamura, A., et al. (2021). Effects of rs958804 and rs7858836 single-nucleotide polymorphisms of the ASTN2 gene on pain-related phenotypes in patients who underwent laparoscopic colectomy and mandibular sagittal split ramus osteotomy. Neuropsychopharmacol Rep 41, 82–90.

Jostins, L., Ripke, S., Weersma, R.K., Duerr, R.H., McGovern, D.P., Hui, K.Y., Lee, J.C., Philip Schumm, L., Sharma, Y., Anderson, C.A., et al. (2012). Host–microbe interactions have shaped the genetic architecture of inflammatory bowel disease. Nature 491, 119–124.

Jung, I., Schmitt, A., Diao, Y., Lee, A.J., Liu, T., Yang, D., Tan, C., Eom, J., Chan, M., Chee, S., et al. (2019). A compendium of promoter-centered long-range chromatin interactions in the human genome. Nat. Genet. 51, 1442–1449.

Koob, G.F., and Volkow, N.D. (2010). Neurocircuitry of addiction. Neuropsychopharmacology 35, 217–238.

Kuleshov, M.V., Jones, M.R., Rouillard, A.D., Fernandez, N.F., Duan, Q., Wang, Z., Koplev, S., Jenkins, S.L., Jagodnik, K.M., Lachmann, A., et al. (2016). Enrichr: a comprehensive gene set enrichment analysis web server 2016 update. Nucleic Acids Res. 44, W90–W97.

Leek, J.T., Johnson, W.E., Parker, H.S., Jaffe, A.E., and Storey, J.D. (2012). The sva package for removing batch effects and other unwanted variation in high-throughput experiments. Bioinformatics 28, 882–883.

Levey, D.F., Gelernter, J., Polimanti, R., Zhou, H., Cheng, Z., Aslan, M., Quaden, R., Concato, J., Radhakrishnan, K., Bryois, J., et al. (2020). Reproducible Genetic Risk Loci for Anxiety: Results From ∼200,000 Participants in the Million Veteran Program. AJP 177, 223–232.

Levis, S.C., Bentzley, B.S., Molet, J., Bolton, J.L., Perrone, C.R., Baram, T.Z., and Mahler, S.V. (2019). On the early life origins of vulnerability to opioid addiction. Mol. Psychiatry.

Leza, L., Siria, S., López-Goñi, J.J., and Fernández-Montalvo, J. (2021). Adverse childhood experiences (ACEs) and substance use disorder (SUD): A scoping review. Drug Alcohol Depend. 221, 108563.

Li, H. (2013). Aligning sequence reads, clone sequences and assembly contigs with BWA-MEM.

Li, H., Handsaker, B., Wysoker, A., Fennell, T., Ruan, J., Homer, N., Marth, G., Abecasis, G., Durbin, R., and 1000 Genome Project Data Processing Subgroup (2009). The Sequence Alignment/Map format and SAMtools. Bioinformatics 25, 2078–2079.

Linnér, R.K., Biroli, P., Kong, E., Meddens, S.F.W., Wedow, R., Fontana, M.A., Lebreton, M., Tino, S.P., Abdellaoui, A., Hammerschlag, A.R., et al. (2019). Genome-wide association analyses of risk tolerance and risky behaviors in over 1 million individuals identify hundreds of loci and shared genetic influences. Nat. Genet. 51, 245–257.

Liu, M., Jiang, Y., Wedow, R., Li, Y., Brazel, D.M., Chen, F., Datta, G., Davila-Velderrain, J., McGuire, D., Tian, C., et al. (2019). Association studies of up to 1.2 million individuals yield new insights into the genetic etiology of tobacco and alcohol use. Nat. Genet. 51, 237–244.

Love, M.I., Huber, W., and Anders, S. (2014). Moderated estimation of fold change and dispersion for RNA-seq data with DESeq2. Genome Biol. 15, 550.

Love, M.I., Hogenesch, J.B., and Irizarry, R.A. (2016). Modeling of RNA-seq fragment sequence bias reduces systematic errors in transcript abundance estimation. Nat. Biotechnol. 34, 1287–1291.

Lundberg, S.M., Erion, G., Chen, H., DeGrave, A., Prutkin, J.M., Nair, B., Katz, R., Himmelfarb, J., Bansal, N., and Lee, S.-I. (2020). From Local Explanations to Global Understanding with Explainable AI for Trees. Nat Mach Intell 2, 56–67.

Manning, A.K., Hivert, M.-F., Scott, R.A., Grimsby, J.L., Bouatia-Naji, N., Chen, H., Rybin, D., Liu, C.-T., Bielak, L.F., Prokopenko, I., et al. (2012). A genome-wide approach accounting for body mass index identifies genetic variants influencing fasting glycemic traits and insulin resistance. Nat. Genet. 44, 659–669.

Martin, M. (2011). Cutadapt removes adapter sequences from high-throughput sequencing reads. EMBnet.journal 17, 10–12.

Maurano, M.T., Humbert, R., Rynes, E., Thurman, R.E., Haugen, E., Wang, H., Reynolds, A.P., Sandstrom, R., Qu, H., Brody, J., et al. (2012). Systematic Localization of Common Disease-Associated Variation in Regulatory DNA. Science 337, 1190–1195.

McInnes, L., Healy, J., and Melville, J. (2018). UMAP: Uniform Manifold Approximation and Projection for Dimension Reduction.

Montez, J.K., and Hayward, M.D. (2014). Cumulative childhood adversity, educational attainment, and active life expectancy among U.S. adults. Demography 51, 413–435.

Mootha, V.K., Lindgren, C.M., Eriksson, K.-F., Subramanian, A., Sihag, S., Lehar, J., Puigserver, P., Carlsson, E., Ridderstråle, M., Laurila, E., et al. (2003). PGC-1α-responsive genes involved in oxidative phosphorylation are coordinately downregulated in human diabetes. Nat. Genet. 34, 267–273.

Morrow, J.J., Bayles, I., Funnell, A.P.W., Miller, T.E., Saiakhova, A., Lizardo, M.M., Bartels, C.F., Kapteijn, M.Y., Hung, S., Mendoza, A., et al. (2018). Positively selected enhancer elements endow osteosarcoma cells with metastatic competence. Nat. Med. 24, 176–185.

Murthy, S., and Gould, E. (2020). How Early Life Adversity Influences Defensive Circuitry. Trends Neurosci. 43, 200–212.

Okbay, A., Baselmans, B.M.L., De Neve, J.-E., Turley, P., Nivard, M.G., Fontana, M.A., Meddens, S.F.W., Linnér, R.K., Rietveld, C.A., Derringer, J., et al. (2016a). Genetic variants associated with subjective well-being, depressive symptoms, and neuroticism identified through genome-wide analyses. Nat. Genet. 48, 624–633.

Okbay, A., Beauchamp, J.P., Fontana, M.A., Lee, J.J., Pers, T.H., Rietveld, C.A., Turley, P., Chen, G.-B., Emilsson, V., Meddens, S.F.W., et al. (2016b). Genome-wide association study identifies 74 loci associated with educational attainment. Nature 533, 539–542.

Patro, R., Duggal, G., Love, M.I., Irizarry, R.A., and Kingsford, C. (2017). Salmon provides fast and bias-aware quantification of transcript expression. Nat. Methods 14, 417–419.

Qiao, X., Zhu, Y., Dang, W., Wang, R., Sun, M., Chen, Y., Shi, Y., and Zhang, L. (2021). Dual-specificity phosphatase 15 (DUSP15) in the nucleus accumbens is a novel negative regulator of morphine-associated contextual memory. Addict. Biol. 26, e12884.

Ramírez, F., Ryan, D.P., Grüning, B., Bhardwaj, V., Kilpert, F., Richter, A.S., Heyne, S., Dündar, F., and Manke, T. (2016). deepTools2: a next generation web server for deep-sequencing data analysis. Nucleic Acids Res. 44, W160–W165.

Ribeiro, A.F., Correia, D., Torres, A.A., Boas, G.R.V., Rueda, A.V.L., Camarini, R., Chiavegatto, S., Boerngen-Lacerda, R., and Brunialti-Godard, A.L. (2012). A transcriptional study in mice with different ethanol-drinking profiles: possible involvement of the GABA(B) receptor. Pharmacol. Biochem. Behav. 102, 224–232.

Rietveld, C.A., Medland, S.E., Derringer, J., Yang, J., Esko, T., Martin, N.W., Westra, H.-J., Shakhbazov, K., Abdellaoui, A., Agrawal, A., et al. (2013). GWAS of 126,559 Individuals Identifies Genetic Variants Associated with Educational Attainment. Science 340, 1467–1471.

Sallari, R.C., Sinnott-Armstrong, N.A., French, J.D., Kron, K.J., Ho, J., Moore, J.H., Stambolic, V., Edwards, S.L., Lupien, M., and Kellis, M. (2017). Convergence of dispersed regulatory mutations predicts driver genes in prostate cancer. Biorxrv.

Schizophrenia Working Group of the Psychiatric Genomics Consortium (2014). Biological insights from 108 schizophrenia-associated genetic loci. Nature 511, 421–427.

Soneson, C., Love, M.I., and Robinson, M.D. (2015). Differential analyses for RNA-seq: transcript-level estimates improve gene-level inferences. F1000Res. 4.

Speliotes, E.K., Willer, C.J., Berndt, S.I., Monda, K.L., Thorleifsson, G., Jackson, A.U., Lango Allen, H., Lindgren, C.M., Luan, J. ‘an, Mägi, R., et al. (2010). Association analyses of 249,796 individuals reveal 18 new loci associated with body mass index. Nat. Genet. 42, 937–948.

Srivastava, A., Malik, L., Sarkar, H., Zakeri, M., Almodaresi, F., Soneson, C., Love, M.I., Kingsford, C., and Patro, R. (2020). Alignment and mapping methodology influence transcript abundance estimation. Genome Biol. 21, 239.

Stahl, E.A., Breen, G., Forstner, A.J., McQuillin, A., Ripke, S., Trubetskoy, V., Mattheisen, M., Wang, Y., Coleman, J.R.I., Gaspar, H.A., et al. (2019). Genome-wide association study identifies 30 loci associated with bipolar disorder. Nat. Genet. 51, 793–803.

Subramanian, A., Tamayo, P., Mootha, V.K., Mukherjee, S., Ebert, B.L., Gillette, M.A., Paulovich, A., Pomeroy, S.L., Golub, T.R., Lander, E.S., et al. (2005). Gene set enrichment analysis: a knowledge-based approach for interpreting genome-wide expression profiles. Proc. Natl. Acad. Sci. U. S. A. 102, 15545–15550.

Sullivan, P.F., Agrawal, A., Bulik, C.M., Andreassen, O.A., Børglum, A.D., Breen, G., Cichon, S., Edenberg, H.J., Faraone, S.V., Gelernter, J., et al. (2018). Psychiatric Genomics: An Update and an Agenda. AJP 175, 15–27.

Teslovich, T.M., Musunuru, K., Smith, A.V., Edmondson, A.C., Stylianou, I.M., Koseki, M., Pirruccello, J.P., Ripatti, S., Chasman, D.I., Willer, C.J., et al. (2010). Biological, clinical and population relevance of 95 loci for blood lipids. Nature 466, 707–713.

Thorvaldsdóttir, H., Robinson, J.T., and Mesirov, J.P. (2013). Integrative Genomics Viewer (IGV): high-performance genomics data visualization and exploration. Brief. Bioinform. 14, 178–192.

Trynka, G., Sandor, C., Han, B., Xu, H., Stranger, B.E., Liu, X.S., and Raychaudhuri, S. (2013). Chromatin marks identify critical cell types for fine mapping complex trait variants. Nat. Genet. 45, 124–130.

Vaiserman, A.M., and Koliada, A.K. (2017). Early-life adversity and long-term neurobehavioral outcomes: epigenome as a bridge? Hum. Genomics 11, 34.

Watanabe, K., Taskesen, E., van Bochoven, A., and Posthuma, D. (2017). Functional mapping and annotation of genetic associations with FUMA. Nat. Commun. 8, 1826.

Widom, C.S., Marmorstein, N.R., and White, H.R. (2006). Childhood victimization and illicit drug use in middle adulthood. Psychol. Addict. Behav. 20, 394–403.

Wolen, A.R., Phillips, C.A., Langston, M.A., Putman, A.H., Vorster, P.J., Bruce, N.A., York, T.P., Williams, R.W., and Miles, M.F. (2012). Genetic dissection of acute ethanol responsive gene networks in prefrontal cortex: functional and mechanistic implications. PLoS One 7, e33575.

Zhang, Y., Liu, T., Meyer, C.A., Eeckhoute, J., Johnson, D.S., Bernstein, B.E., Nusbaum, C., Myers, R.M., Brown, M., Li, W., et al. (2008). Model-based analysis of ChIP-Seq (MACS). Genome Biol. 9, R137.

Zhou, H., Rentsch, C.T., Cheng, Z., Kember, R.L., Nunez, Y.Z., Sherva, R.M., Tate, J.P., Dao, C., Xu, K., Polimanti, R., et al. (2020a). Association of OPRM1 Functional Coding Variant With Opioid Use Disorder: A Genome-Wide Association Study. JAMA Psychiatry.

Zhou, H., Sealock, J.M., Sanchez-Roige, S., Clarke, T.-K., Levey, D.F., Cheng, Z., Li, B., Polimanti, R., Kember, R.L., Smith, R.V., et al. (2020b). Genome-wide meta-analysis of problematic alcohol use in 435,563 individuals yields insights into biology and relationships with other traits. Nat. Neurosci. 23, 809–818.

